# Supervised machine learning to decipher the complex associations between neuro-immune biomarkers and quality of life in schizophrenia

**DOI:** 10.1101/393942

**Authors:** Buranee Kanchanatawan, Michael Maes

**Affiliations:** Department of Psychiatry, Faculty of Medicine, Chulalongkorn University, Patumwan, Bangkok, 10330, Thailand; Department of Psychiatry, Medical University of Plovdiv, Plovdiv, 4000, Bulgaria; IMPACT Strategic Research Center, Deakin University, PO Box 281, Geelong, 3220, Australia

**Keywords:** immune, inflammation, tryptophan catabolites, deficit schizophrenia, depression, physio-somatic

## Abstract

Stable phase schizophrenia is characterized by altered patterning in tryptophan catabolites (TRYCATs) and memory impairments, which are associated with PHEMN (psychosis, hostility, excitation, mannerism and negative) and DAPS (depression, anxiety and physio-somatic) symptoms. This study was carried out to examine the association between TRYCAT patterning, memory impairments, psychopathological features and health-related quality of life (HR-QoL) in schizophrenia.

The World Health Organization (WHO) QoL instrument-Abbreviated version (WHO-QoL-BREF), IgA/IgM responses to TRYCATs, cognitive tests, Scale for the Assessment of Negative Symptoms (SANS), Hamilton and Depression (HAMD) and Anxiety (HAMA) Rating Scales and the Fibromyalgia and Chronic Fatigue Syndrome Rating Scale (FF) were measured in 80 schizophrenia patients and 40 controls.

Neural Network analysis shows that the total HR-Qol score is best predicted by (in descending order) FF, HAMA and SANS scores, Mini Mental State examination, hostility, ratio noxious/protective TRYCATs and HAMD score. Partial least Squares (PLS) analysis shows that 55.1% of the variance in Domain1 (physical) is predicted by PHEMN and DAPS latent vector (LV) scores, while 57.9% of domain2 (psychological), 32.7% of domain3 (social) and 55.0% of domain4 (environment) are explained by DAPS LV scores. TRYCATs and episodic/semantic memory impairments have specific indirect effects on domains 2, 3 and 4, which are mediated by DAPS symptoms, while the effects of TRYCATs on domain1 are mediated by PHEMN and DAPS symptoms. Picolinic acid, xanthurenic acid and 3-hydroxy-kynurenine decrease WHO-QoL scores, whilst anthranilic acid is protective.

The results show that lowered HR-Qol in schizophrenia is strongly predicted by noxious TRYCATs, impairments in episodic and semantic memory and DAPS symptoms, especially physio-somatic symptoms and anxiety. Neuro-immune pathways and the consequent cognitive impairments determine to a great extent lowered HR-QoL in schizophrenia.

## Introduction

In 1995, Smith and Maes (1995) launched the monocyte-T lymphocyte theory of schizophrenia considering that activated immune-inflammatory pathways explain the neurodevelopmental pathology linked with infections in utero through the adverse effects of activated immune-inflammatory pathways, activation of microglia and the tryptophan catabolite (TRYCATs) pathway, oxidative and nitrosative stress (O&NS), and modulation of glutamate production and N-methyl D-aspartate (NMDA) receptor function. Infectious hits occurring in utero combined with secondary hits during adolescence may unveil positive and negative symptoms of schizophrenia in susceptible individuals (Anderson et al., 2013a; 2013b; Davis et al., 2014; 2016). The ensuing neurotoxic effects of activated immune-inflammatory, O&NS and TRYCAT pathways coupled with epigenetic changes may cause early maturation disorders in developing neuronal brain structures, while later hits may lead to neuroprogression, that is the process of progressing brain neuro-circuitry abnormalities caused by neuro-immune, neuro-oxidative and neuro-nitrosative pathways including neuropil shrinkage, and disorders in neural connectivity, neuronal dendrite growth, neuronal plasticity, neurogenesis, neuronal apoptosis and membrane protein expression (Anderson et al., 2013a; 2013b; Davis et al., 2014; 2016).

There is evidence that acute and stable phases of schizophrenia are accompanied by activated immune-inflammatory pathways including elevated levels of M1 macrophagic, T helper (Th)-1-like, Th-2-like and Tregulatory (Treg) cytokines and by enhanced oxidative stress pathways (Maes et al., 1994; 1995a; 1995b; Noto et al., 2015a; 2015b; 2016; Brinholi et al., 2015). One consequence of activation of these pathways is stimulation of indoleamine 2,3-dioxygenase (IDO), which lowers plasma tryptophan and increases noxious TRYCATs, including picolinic acid (PA), xanthurenic acid (XA), kynurenine (KYN), 3-OH-kynurenine (3HK) and quinolinic acid (QA), and generally more protective TRYCATs, including anthranilic acid (AA) and kynurenic acid (KA) (Kanchanatawan et al., 2018a; 2018c). Recently we discovered that schizophrenia, and especially deficit schizophrenia, is accompanied by increased levels of IgA responses directed against PA, XA, 3-HK and QA and an increased noxious / protective TRYCAT ratio, indicating increased impact of noxious TRYCATs (Kanchanatawan et al., 2018c). In addition, we found that the IgM responses to noxious TRYCATs were significantly lower in patients with deficit versus nondeficit schizophrenia, suggesting lowered regulation of noxious TRYCATs (Kanchanatawan et al., 2018b). This is important as PA, XA, 3-HK and QA have excitotoxic, cytotoxic, inflammatory and oxidative effects. Moreover, elevations in peripheral TRYCAT concentrations such as kynurenine, QA and 3-HK determine to a large extent part brain QA concentrations (Kita et al., 2002).

Recently, we observed that a) stable phase schizophrenia is characterized by two interrelated symptom dimensions, a first comprising psychosis, hostility, excitation, mannerism and negative (PHEMN) symptoms, and a second depressive, anxiety and physio-somatic (DAPS) symptoms (Kanchanatawan et al., 2018e); and 2) a large part of the variance in PHEMN and DAPS factor scores was explained by changes in TRYCAT patterning and cognitive deficits, including in episodic and semantic memory, as assessed with Consortium to Establish a Registry for Alzheimer’s disease (CERAD) tests (Kanchanatawan et al., 2018a). Moreover, we found that the combined use of TRYCAT levels and cognitive features strongly segregates deficit from nondeficit schizophrenia and healthy controls, with TRYCAT levels and false memory recall being the most important features of deficit schizophrenia (Kanchanatawan et al., 2018d).

There is now consensus that the assessment of psychopathological symptoms, including negative and positive symptoms, is insufficient to evaluate the intermediate and distal outcome of schizophrenia (Galuppi et al., 2010; Bobes et al., 2007). Health-related quality of life (HR-QoL) and social functioning are now firmly consolidated distal outcome measurements, which are broadly demanded by families, patients and clinicians (Galuppi et al., 2010; Bobes et al., 2007). Unfortunately, the precise factors which impact HR-QoL in schizophrenia are not well established, although studies reported the impact of negative, positive and affective symptoms (Sim et al., 2004; Ritsner et al., 2005). For example, depression and anxiety, but not negative or positive, symptoms predict HR-Qol (Becker et al., 2005; Fitzgerald et al., 2001), while other studies showed that negative and positive symptoms are associated with HR-QoL in schizophrenia (Norman et al., 2000). These discrepancies may in part be explained by our recent findings that in schizophrenia mood, negative and positive symptoms are strongly interrelated and that positive symptoms should be further “dissected” into PHEM symptoms (Kanchanatwan et al., 2018e). Moreover, there are no data whether the neuro-immune and cognitive features of schizophrenia may impact HR-QoL.

Hence, the aim of this study is to examine the associations between TRYCAT levels and HR-QoL in stable phase schizophrenia considering the possible causal effects of TRYCATs, cognitive impairments (using the CERAD), and DAPS/PHEMN symptoms on HR-QoL. Previous research examined associations between HR-QoL and negative/positive/mood symptoms using correlational or regression analyses. Multiple Linear Regression Analysis (MLRA) is adequate when there a few explanatory variables, which are not significantly collinear and have a clear relationship with the outcome variable (Tobias, 1995). However, changes in TRYCAT patterning, episodic and semantic memory tests and DAPS/PHEMN symptoms are strongly interrelated and therefore MLRA is less adequate. In order to examine multiple interrelated factors, two different supervised machine learning techniques are advocated, namely neural network (NN) and partial least squares (PLS) analyses (Tobias, 1995; Land and Rettenmeier, 2017; Kitikidou and Iliadis, 2012; Maitra and Yan, 2008). NN is well suited to discover more complex nonlinear relationships between input variables (TRYCATs, cognitive tests and symptomatology) predicting HR-QoL (the outcome) and additionally allows to rank the input variables in order of predictive relevance (Kitikidou and Iliadis, 2012). PLS extracts relevant latent factors, which account for as much of the variance as possible in the input variables while being relevant to the latent factor extracted from the response (HR-QoL) variable (Tobias, 1995).

## Subjects and Methods

### Participants

In this case-control study we included Thai adults, 18 to 65 years, namely outpatients with stable phase schizophrenia and normal controls. Participants with schizophrenia were recruited at the Department of Psychiatry, Chulalongkorn University, Bangkok, Thailand. Patients with schizophrenia were diagnosed using DSM-IV-TR and DSM-5 criteria by the principal investigator (BK). Towards this end the authors used the Thai validated version of the Mini*-*International Neuropsychiatric Interview (M.I.N.I.) (Kittirathanapaiboon and Khamwongpin, 2005). Moreover, we made the diagnosis of primary deficit schizophrenia using the Schedule for the Deficit Syndrome (SDS) (Kirkpatrick et al., 1989). Schizophrenia patients who did not comply with SDS criteria of deficit schizophrenia were classified as nondeficit schizophrenia. Healthy individuals were recruited by word of mouth from the same catchment area, Bangkok, Thailand. The following schizophrenia patients were excluded: participants with a lifetime diagnosis of axis-I DSM-IV-TR / DSM-5 disorders, such as major depression, bipolar disorder, psycho-organic syndrome and substance use disorders. We excluded normal controls with a lifetime / current diagnosis of axis-I DSM-IV-TR / DSM-5 disorders. We also excluded participants with medical illness, including inflammatory bowel disease, rheumatoid arthritis, chronic obstructive pulmonary disease, multiple sclerosis, diabetes type 2, neurodegenerative disorders and subjects who underwent any treatment with immunomodulatory drugs and antioxidant supplements. All participants (controls and patients) as well as guardians of all patients gave written informed consent prior to participation in this study. The study was conducted according to Thai and international ethics and privacy laws. Approval for the study was obtained from the Institutional Review Board of the Faculty of Medicine, Chulalongkorn University, Bangkok.

### 2.2 Procedures

All participants underwent a comprehensive semistructured interview performed by BK who scored all rating scales, made the diagnoses and applied inclusion and exclusion criteria. We used the M.I.N.I. in a validated Thai translation (Kittirathanapaiboon and Khamwongpin, 2005), the total score on Scale for the Assessment of Negative Symptoms (SANS) (Andreasen, 1989), the postive and negative symptom subscales of the Positive and Negative Syndrome Scale (PANSS) (Kay et al., 1986), and the Schedule for Deficit Syndrome (SDS) (Kirkpatrick et al., 1989). Moreover, the Brief Psychiatric Rating Scale (Overall and Gorham, 1962) was used to assess psychopahthology. Beside using the SANS, we used BPRS and PANSS items to compute the 4 remaining PHEMN dimensions (Kanchanatawan et al., 2018e). Psychosis was assessed as a z unit weighted composite score computed as z score PANSS P1 (delusion) (zP1) + zP3 (hallucinations) + zP6 (suspiciousness) + zBPRS11 (suspiciousness) + zBPRS12 (hallucinatory behavior) + BPRS15 (unusual thought content). The Hostility dimension was assessed as zP7 (hostility) + zPANSS general14 (zG14, poor impulse control) + zBPRS10 (hostility) + zBPRS14 (uncooperativeness). The excitement dimension was computed as zP14 (excitement) + zP5 (grandiosity) + zBPRS8 (grandiosity) + zBPRS17 (excitement). The mannerism-posturing dimension was assessed as zG5 + zBPRS7 (mannerism and posturing). Severity of depression was measured using the Hamilton Depression (HAMD) Rating Scale, while anxiety was scored using the Hamilton Anxiety (HAM-A) Rating Scale (Hamilton, 1959; 1960). Severity of physio-somatic symptoms was assessed employing the 12 item Fibromyalgia and Chronic Fatigue Syndrome Rating scale (FF) (Zachrisson et al., 2002). We made the diagnosis of nicotine dependence according to DSM-IV-TR criteria. Moreover, socio-demographic data were assessed, including years of education, employment and marital status, number of lifetime psychotic episodes and use of psychopharmacological drugs. Body mass index (BMI) was computed as weight (in kg) divided by square of height (in m^2^).

The same day as the semistructured interview we measured HR-QoL employing the World Health Organization Quality of Life instrument-Abbreviated version (WHO-QoL-BREF) (WHO, 1993). This scale scores 26 items measuring four HR-QoL domains: 1) Domain 1 or physical health: energy, sleep, fatigue, pain, discomfort, work capacity, activities of daily living, dependence on medicinal substances, mobility; 2) Domain 2 or psychological health: self-esteem, bodily image, learning, thinking, concentration, memory, negative and positive feelings, beliefs (spirituality-religion-personal); 3) Domain 3 or social relationships: social support, sexual activity, personal relationships; and 4) Domain 4 or environment: physical safety and security, health and social care, freedom, financial resources, home environment, recreation and leisure activities and physical environment. We computed raw scores on the 4 domains according to the WHO-QoL-BREF criteria (WHO, 1993) and computed the total WHO-QoL-BREF score by summing up the 26 item scores.

The same day as the semistructured interview, a clinical research assistant (blinded to the clinical diagnosis) with a master degree in mental health scored 5 CERAD tests (Consortium to Establish a Registry for Alzheimer’s disease) (CERAD, 1986; Welsh et al., 1994): Mini-Mental State Examination (MMSE) to measure orientation, concentration, memory, speech and constructional praxis; Verbal Fluency Test (VFT) to measure fluency or semantic memory; Word List Memory (WLM) to assess verbal episodic memory and immediate working memory; Word list recall - False Recall to measure false memory creation; and Word List Recognition (WLRecog) to measure verbal learning or recall recognition.

### Biomarkers

At 8.00 a.m. fasting blood was sampled the same day we completed interviews and CERAD tests. Blood was frozen at -80 °C until thawed for assay of IgA and IgM responses directed to QA, 3HK, PA, XA, KA and AA (Kanchanatawan et al., 2018b; 2018c). Briefly, “The 6 TRYCATs were dissolved in 200 μL dimethylsulfoxide (DMSO) (Acros). Bovine serum albumin (BSA) (ID Bio) was dissolved in 3mL 2-morpholino-ethanesulfonic acid monohydrate (MES Acros) buffer 10-1 M at pH = 6.3 (Acros). The TRYCATs were then mixed with the BSA solution and supplemented with 15 mg N-hydroxysuccinimide (Sigma) and 1-(3-dimethylaminopropyl)-3-ethylcarbodiimide (Acros) as coupling agents. The conjugates were synthesized by linking 3HK (Sigma), KA (Acros), QA (Acros), AA (Acros), XA (Acros) and PA (Acros) to 20 mg BSA. The coupling reaction proceeded at 37°C for 1 hour in the dark. The coupling was stopped by adding 100 mg hydroxylamine (Sigma-Aldrich) per conjugate. Protein conjugates were dialyzed with 10-1 M NaCl solution, pH=6 for 72 hours, with the bath solution being changed at least four times per day. The conjugated TRYCATs and BSA concentrations were evaluated by spectrophotometry. The coupling ratio of each conjugate was determined by measuring the concentration of TRYCATs and BSA at 310–330 nm and 280 nm, respectively. ELISA tests were used to determine plasma titers of serum immunoglobulin (Ig)M and IgA. Towards this end, polystyrene 96-well plates (NUNC) were coated with 200 μL solution containing 10–50 μg/mL TRYCAT conjugates in 0.05 M carbonate buffer (pH = 9.6). Well plates were incubated under agitation at 4 °C for 16 hours. Then, 200 μL blocking buffer A (Phosphate Buffered Saline, PBS, 2.5 g/L BSA, pH=7) was applied and all samples were incubated at 37 °C for 1 hour. Well plates were washed with PBS solution and filled with 100 μL serum diluted 1:130 in blocking buffer and incubated at 37 °C for 1 hour and 45 minutes. Well plates were washed 3 times with PBS, 0.05% Tween 20, incubated with peroxidase-labeled goat anti-human IgA (SouthernBiotech) antibodies at 37 °C for 1 hour. The goat anti-human IgM antibody was diluted at 1:5000 and the IgA antibody was diluted at 1:10,000 in blocking buffer (PBS, 2.5 g/L BSA). Plates were then washed three times with PBS, 0.05% Tween 20. Fifty µL of 3,3’,5,5’-Tetramethylbenzidine (TMB) substrate (SouthernBiotech) was added and incubated for 10 minutes in the dark. The reaction was stopped using 50 µl of TMB stop solution (SouthernBiotech). Optical densities (ODs) were measured at 450 nm using Varioskan Flash (Thermo Scientific). All assays were carried out in duplicate. The analytical intra-assays CV values were < 7%. The OD scores were expressed as z-united weighted scores”. We computed 3 ratio as explained previously (Kanchanatawan et al., 2018b; 2018c): a) zIgA NOX_PRO = sum of z scores of QA (zQA) + zPA + zXA – zAA – zKA (index of increased noxious potential); b) ΔNOX_PRO = zIgA (zQA + zPA + zXA +z3HK – zAA – zKA) – zIgM (zQA + zPA + zXA +3HK – zAA – zKA) (a more comprehensive index of increased noxious potential); and c_ zIgM KA_3HK = zIgM KA – z3HK (index of lowered regulation of KA versus 3HK).

### Statistical analyses

Intergroup differences in demographic and clinical dimensions were assessed employing analyses of variance (ANOVAs), while associations among categorical variables were checked using analyses of contingency tables (χ^2^ tests). We divided the participants into three equal groups based on the sum of the 4 domains giving very low (<74.6667, n=39), low (between 74.6667 and 92.3333, n=40) and high (> 92.3333, n=39) WHO-QoL-BREF scores. P-corrections for false discovery rate were used to assess the results of multiple statistical analyses (Benjamini and Hochberg, 1995). We used multivariate GLM analyses to assess the effects of explanatory variables on the WHOQoL-BREF domain and total scores. Consequently, tests for between-subject effects were used to analyze the associations between significant explanatory and dependent variables. Model-generated estimated marginal mean values were interpreted using protected, pair-wise post-hoc analyses to examine intergroup differences. We used linear multiple regression analyses to examine the associations between one dependent variable (total or domain WHO-QoL scores) and multiple explanatory variables and used partial eta squared values to estimate effects sizes. Results of regression analyses were checked for multicollinearity. We used the IBM SPSS Windows version 24 to analyze data. Statistical significance was set at 0.05, two-tailed.

In this study, the authors employed multilayer perceptron (MLP) Neural Network (NN) analyses (SPSS 24) to examine the more complex associations between the total WHO-QoL score (output variable) and TRYCATs, CERAD tests and PHEMN/DAPS items (with age, sex and education) as input variables in an automated feedforward architecture model. The network was trained using one or two hidden layers with a variable number of nodes. The percentage of cases assigned to the training (to estimate the network parameters), testing (to prevent overtraining) and holdout (to evaluate the final network) sets were 46.67%, 20.0% and 33.33%, respectively. The stopping rule was one consecutive step with no further decrease in the error term. Error, relative error and (relative) importance of each of the input variables were computed in sensitivity analyses.

In order to examine the causal links among the TRYCAT, cognitive, symptoms and WHO-QoL variables we used Partial Least Squares (PLS) analysis using SmartPLS software (Ringle et al., 2015). PLS is a structural equation modeling technique which employs pathway modeling performed on latent vectors extracted from indicator variables coupled with a PLS-structural equation modeling algorithm (Ringle et al., 2014). We entered WHO-QoL data as final output or response variable, e.g. by entering the 4 domain scores as indicator variables (in a reflective model) and then extracting the latent construct “WHO-QoL”. All 8 DAPS/PHEMN symptoms scores were entered as indicators of the latent construct “general psychopathology” or we entered the 3 DAPS and 5 PHEMN scores as latent constructs of the DAPS and PHEMN dimensions, respectively. The 5 CERAD tests were entered as indicators of the latent construct “cognitive functions” and the 3 TRYCAT ratios as indicators of “TRYCAT patterning”. Symptoms and WHO-QoL scores were entered as single indicators (e.g. all 4 domains) or as latent variables as measured with indicator variables in a reflective model. TRYCATs, CERAD and DAPS/PHEMN were entered as direct input variables predicting WHO-QoL scores. CERAD and TRYCAT latent vector (LV) scores were entered as direct predictors of the symptom dimensions, while the TRYCAT LV predicted CERAD tests. Quality of the model was assessed using overall model fit with as criterion a SRMR value < 0.08). In addition, we only accepted constructs with a good reliability and discriminant validity as indicated by Cronbach’s alpha values > 0.7, composite reliability > 0.7 and average variance extracted (AVE) > 0.500. Moreover, indicators were only included in latent constructs when their factor loadings were > 0.5 with p values < 0.001, indicating that the convergent validity is adequate. Finally, we checked construct crossvalidated redundancies and communalities (Ringle et al., 2014). We only performed path analysis when the model and constructs complied with these quality criteria. Subsequently, path coefficients with exact p-value, total effects, total indirect and specific indirect effects were computed.

## Results

### Socio-demographic data

**Table 1** shows the socio-demographic data of the participants in this study divided into three groups according to the q33 and q66 values of the sum on the 4 domains, namely those with very low, low and normal WHO-Qol total values. Subjects with lower WHO-QoL values were somewhat older than those with normal WHO-QoL values (after p correction: p=0.048). There were no significant differences in sex ratio, marital status and BMI between the WHO-QoL groups. Education was lower in those with very low WHO-QoL values, while the same group also showed a higher frequency of familial history of psychosis than in those with lower WHO-QoL values. The number of psychotic episodes and patients with schizophrenia were higher in the low WHO-QoL group. **Electronic Supplementary File (ESF) Figure 1** shows that there are significant differences in the 4 domain and total WHO-QoL scores among controls and schizophrenia patients with and without deficit syndrome (F=10.49, df=8/224, p<0.001; partial eta squared=0.273). Domain 1, 2 and 4, and total scores were significantly different between the three groups and significantly decreased from control → nondeficit → deficit. Domain3 score was significantly lower in patients with deficit schizophrenia than in controls (p=0.001), while there were no significant differences between the nondeficit type and either controls (p=0.092) or patients with the deficit subtype (p=0.082).

**Table 1.** Socio-demographic and clinical data in normal controls and schizophrenic patients divided according to the total score on the World Health Organization Quality of Life instrument-Abbreviated version (WHO-QoL-BREF)

| Variables | WHO-QoL<br><74.6667<sup>A</sup><br>n=39 | WHO-QoL 74.6667-92.3333<sup>B</sup><br>n=40 | WHO-QoL > 92.3333<sup>C</sup><br>n=39 | F/X<sup>2</sup> | df | p |
| --- | --- | --- | --- | --- | --- | --- |
| Age (years) | 41.9 (10.5)<sup>C</sup> | 41.9 (11.8)<sup>C</sup> | 35.6 (12.1)<sup>A,B</sup> | 3.87 | 2/115 | 0.024 |
| Sex (M / F) | 20 / 19 | 16 / 24 | 15 / 24 | 1.56 | 2 | 0.458 |
| Single-divorced / married | 35 / 4 | 29 / 9 | 27 / 12 | 5.01 | 2 | 0.082 |
| BMI (kg/m<sup>2</sup>) | 24.2 (4.3) | 25.4 (5.7) | 23.5 (4.4) | 1.53 | 2/110 | 0.221 |
| Employment (No/Yes) | 26 / 13<sup>C</sup> | 16 / 24 | 7 / 32<sup>A</sup> | 19.12 | 2 | <0.001 |
| Education (years) | 11.5 (4.6)<sup>C</sup> | 12.7 (4.2) | 14.7 (4.5)<sup>B</sup> | 5.48 | 2/115 | 0.005 |
| TUD (No/Yes) | 37 / 2 | 36 / 4 | 38 / 1 | NA | - | - |
| NC /SCZ / deficit | 1 / 14 / 24<sup>B,C</sup> | 9 / 19 / 12<sup>A,C</sup> | 30 / 7 / 2<sup>A,B</sup> | 58.64 | 4 | <0.001 |
| Number psychoses | 3.29 (2.43)<sup>C</sup> | 2.57 (3.35)<sup>C</sup> | 0.46 (0.99)<sup>A,B</sup> | 14.03 | 2/111 | <0.001 |
| Familial History Psychosis<br>(No/Yes) | 22 / 17<sup>C</sup> | 29 / 11 | 34 / 5<sup>A</sup> | 9.17 | 2 | 0.010 |

All results are shown as mean (SD)
NA: results of X<sup>2</sup> test are probably invalid as some cells have expected cell counts less than 5
BMI: body mass index; TUD: tobacco use disorder
NC/SCZ/deficit: ratio of normal controls / schizophrenia patients without deficit syndrome / with deficit syndrome

**Figure 1.**
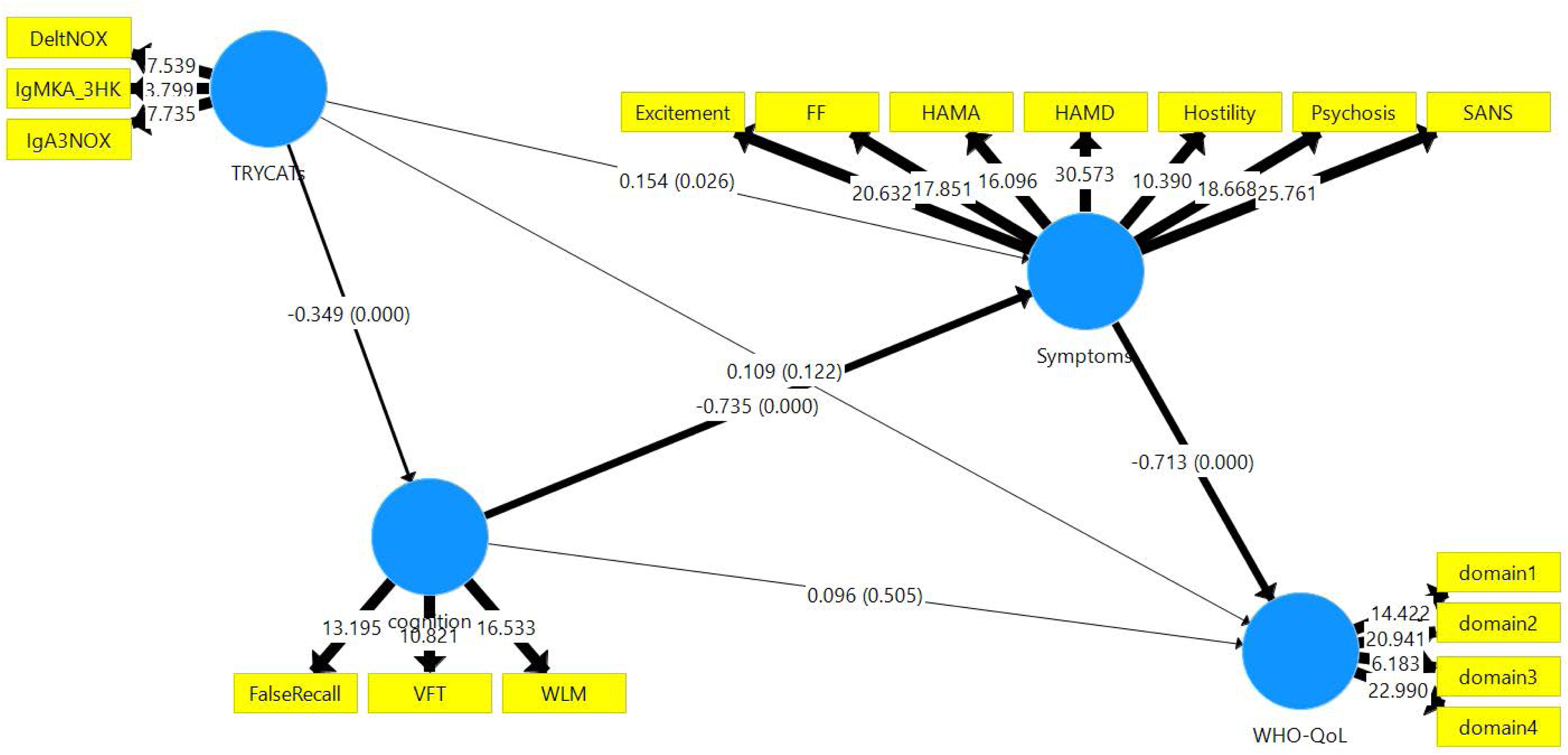
Results of Partial Least Squares path modeling analysis (with path coefficients and exact p-values) with the 4 domains of the WHO-QoL as indicator variables of the latent construct “WHO-QoL”. Input variables are all symptom dimensions, cognitive tests and tryptophan catabolites. See legends to ESF Figure 2, 3 and 4 for explanation

### Differences in clinical ratings between the three WHO-QoL groups

**Table 2** shows the outcome of multivariate GLM analyses with 8 scores (HAMA, HAMD, FF, SANS, Psychosis, Hostility, Excitement and Mannerism) as dependent variables. In Regression #1 we used the three WHO-Qol groups, clinical diagnosis (controls versus deficit versus nondeficit schizophrenia), sex and use of haloperidol as explanatory variables. The other extraneous variables did not reach significance (these variables are listed below). We found that WHO-QoL groups had a significant effect on the 8 rating scales even after adjusting for diagnosis. Regression #2 shows the same multivariate regression but now without diagnosis. We found a strong association between the WHO-QoL groups and the rating scales (explained variance: 34.4%). Tests for between-subject effects showed differences in all 8 scores between the three WHO-QoL subgroups with the strongest associations with FF (39.7%), psychosis (38.9%) and HAMD (36.3% variance). **ESF Figure 2** shows the z-transformed data of the 8 scores in the three study groups, whereas **Table 3** shows the model-generated marginal estimated means obtained by regression #2. Protected post-hoc analyses show that HAMA, HAMD, FF, SANS, Psychosis, Excitement and Mannerism scores are different between the three study groups and increased from the normal WHO-QoL group → low group → very low group. Hostility scores were higher in subjects with the very low WHO-QoL values as compared with the other 2 groups.

**Table 2.** Result of multivariate general linear model (GLM) analysis with the IgA responses directed to tryptophan catabolites (TRYCATS) as dependent variables and groups divided according to the total score on the World Health Organization Quality of Life instrument-Abbreviated version (WHO-QoL-BREF) as primary explanatory variable

| Type of analyses | Dependent Variables | Explanatory Variables | F | df | P | Partial Eta Squared |
| --- | --- | --- | --- | --- | --- | --- |
| **Multivariate #1** | 8 rating scales | WHO-QoL_3groups | 2.37 | 16/204 | 0.003 | 0.157 |
|  |  | HC/SCZ/Deficit | 16.60 | 16/204 | <0.001 | 0.566 |
|  |  | Sex | 2.58 | 8/102 | 0.013 | 0.168 |
|  |  | Haloperidol | 2.94 | 8/102 | 0.005 | 0.187 |
| **Multivariate #2** | 8 rating scales | WHO-QoL_3groups | 6.82 | 16/208 | <0.001 | 0.344 |
|  |  | Sex | 2.32 | 8/104 | 0.025 | 0.152 |
|  |  | Haloperidol | 3.56 | 8/104 | 0.001 | 0.215 |
| **Between subject-effects** | HAMA | WHO-QoL_3groups | 24.64 | 2/111 | <0.001 | 0.307 |
|  | HAMD | WHO-QoL_3groups | 31.65 | 2/111 | <0.001 | 0.363 |
|  | FF | WHO-QoL_3groups | 36.60 | 2/111 | <0.001 | 0.397 |
|  | SANS | WHO-QoL_3groups | 30.94 | 2/111 | <0.001 | 0.358 |
|  | Psychosis | WHO-QoL_3groups | 35.37 | 2/111 | <0.001 | 0.389 |
|  | Hostility | WHO-QoL_3groups | 16.36 | 2/111 | <0.001 | 0.228 |
|  | Excitement | WHO-QoL_3groups | 27.78 | 2/111 | <0.001 | 0.334 |
|  | Mannerism | WHO-QoL_3groups | 14.24 | 2/111 | <0.001 | 0.204 |
| **Multivariate #3** | 3 TRYCAT ratios | WHO-QoL_3groups | 4.05 | 6/222 | 0.001 | 0.099 |
|  |  | Age | 3.18 | 3/111 | 0.027 | 0.079 |
| **Between-subject effects** | IgA NOX_PRO | WHO-QoL_3groups | 9.23 | 2/113 | <0.001 | 0.140 |
|  | Δ NOX_PRO | WHO-QoL_3groups | 3.05 | 2/113 | 0.051 | 0.068 |
|  | IgM KA_3HK | WHO-QoL_3groups | 5.60 | 2/113 | 0.020 | 0.089 |
| **Multivariate #4** | 5 CERAD tests | WHO-QoL_3groups | 4.48 | 10/212 | <0.001 | 0.174 |
|  |  | Sex | 3.38 | 5/106 | 0.007 | 0.137 |
|  |  | Education | 14.54 | 5/106 | <0.001 | 0.407 |
| **Between-subject effects** | WLM | WHO-QoL_3groups | 16.78 | 2/110 | <0.001 | 0.234 |
|  | VFT | WHO-QoL_3groups | 11.16 | 2/110 | <0.001 | 0.169 |
|  | MMSE | WHO-QoL_3groups | 4.01 | 2/110 | 0.021 | 0.068 |
|  | FalseRecall | WHO-QoL_3groups | 9.85 | 2/110 | <0.001 | 0.152 |
|  | WordLRecognition | WHO-QoL_3groups | 2.53 | 2/110 | 0.084 | 0.044 |

WHO-QoL\_3 groups: divided according to WHO-QoL <72.666 (very low), WHO-QoL 72.667-90.333 (low) and WHO-QoL >90.3333 (normal) HAMA/HAMD: Hamilton Anxiety or Depression Rating Scale. FF: the Fibromyalgia and Chronic Fatigue Syndrome Rating scale (FF). SANS: Scale for the Assessment of Negative Symptoms. Psychosis, Hostility, Excitation and Mannerism: computed as z unit weighted composite scores using symptoms of the Brief Psychiatric Rating Scale and the Positive and Negative Syndrome Scale
IgA NOX\_PRO: computed as sum of z scores of QA (zQA) + zPA + zXA – zAA – zKA (index of increased noxious potential); QA: quinolinic acid; PA: picolinic acid; XA: xanthurenic acid; AA: anthranilic acid; KA: kynurenic acid. ΔNOX\_PRO: computed as zIgA (zQA + zPA + zXA + z3HK – zAA – zKA) – zIgM (zQA + zPA + zXA + 3HK – zAA – zKA) (a more comprehensive index of increased noxious potential); 3HK: 3-OH-kynurenine. zIgM KA\_3HK: computed as zIgM KA – z3HK (index of lowered regulation of KA versus 3HK).
CERAD: Consortium to Establish a Registry for Alzheimer's Disease
WLM: Word List Memory
VFT: Verbal Fluency Test
MMSE: Mini mental State Examination

**Table 3.**
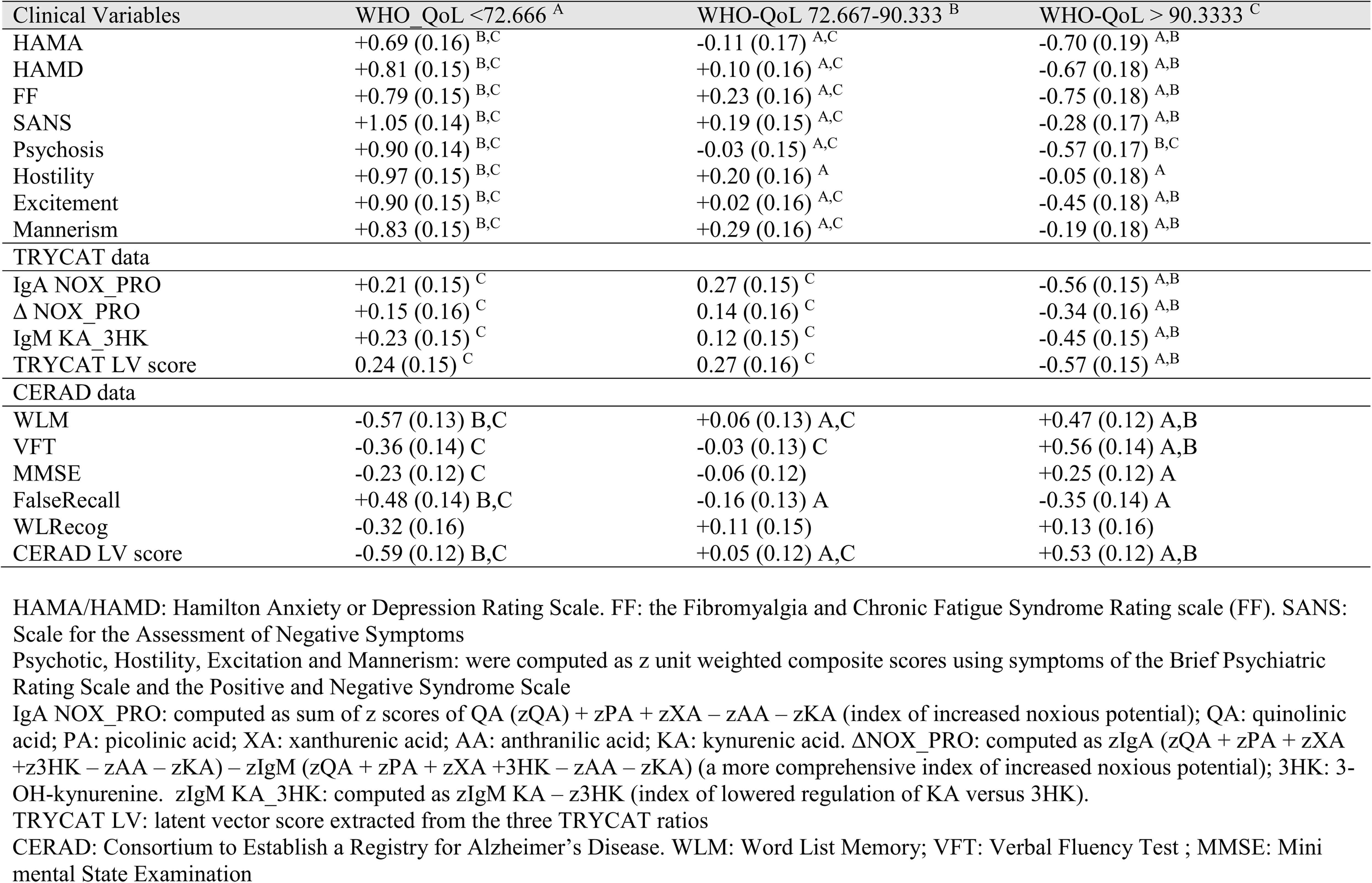
Model-generated estimated marginal mean (SE) values (expressed as z-scores) obtained by multivariate GLM analysis with groups divided according to the total score on the World Health Organization Quality of Life instrument-Abbreviated version (WHO-QoL-BREF) as grouping variable

**Figure 2.**
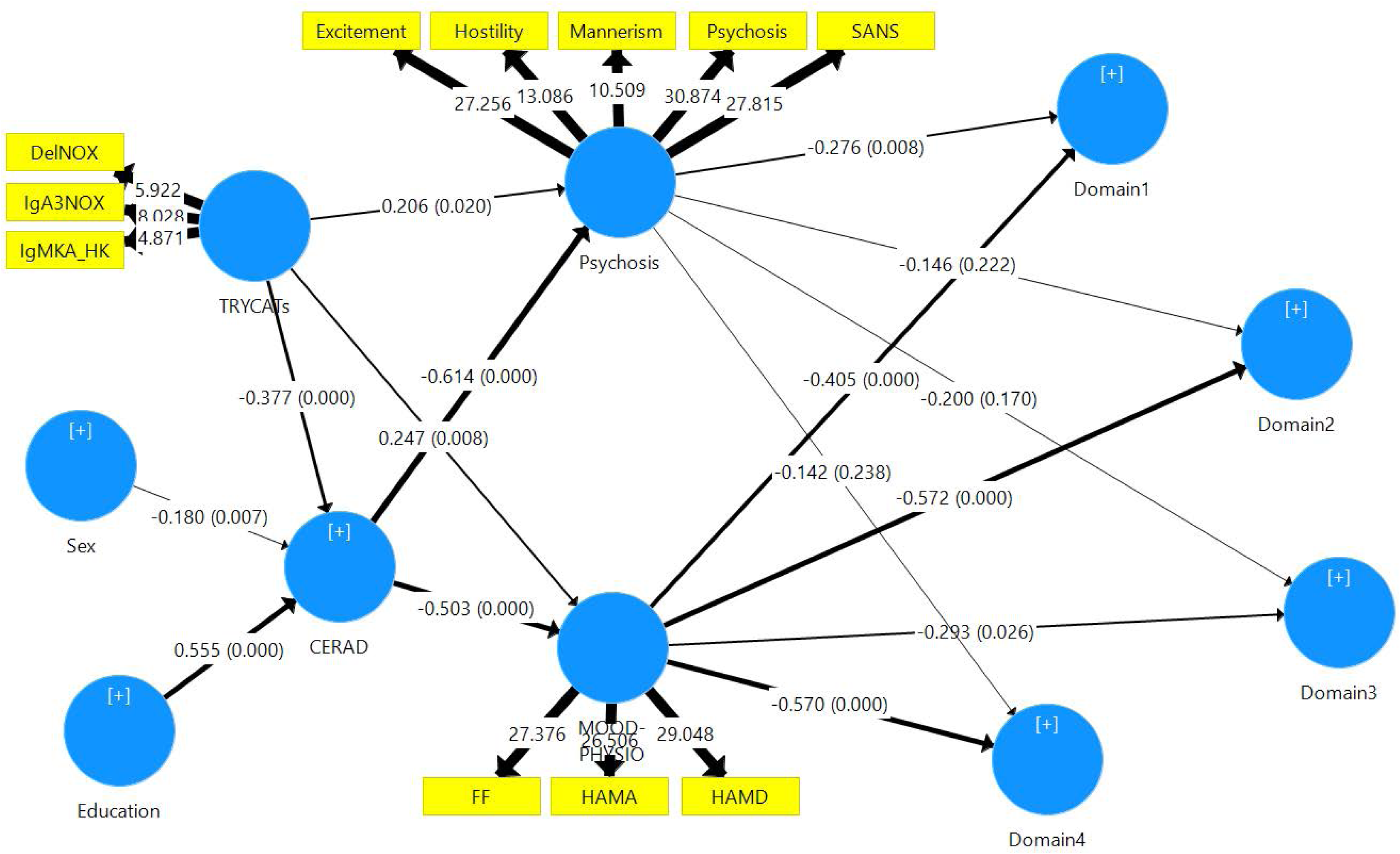
Results of Partial Least Squares path modeling analysis (with path coefficients and exact p-values) with the 4 domains of the WHO-QoL as output (indicator) variables and two symptom dimensions and tryptophan catabolites as input variables. See legends to ESF Figure 2, 3 and 4 for explanation

Multivariate analysis #2 shows also an effect of sex. After p-correction, there were significant effects of sex on SANS (F=4.22, df=1/111, p=0.042), psychosis (F=7.13, df=1/111, p=0.009), hostility (F=14.57, df=1/111, p<0.001), excitation (F=4.82, df=1/111, p=0.030) and mannerism (F=12.63, df=1/111, p=0.001) (all higher in males than females). Multivariate analysis #2 shows also a significant effect of haloperidol being associated with SANS (F=12.26, df=1/111, p=0.001), hostility (F=11.13, df=1/111, p=0.001) and mannerism (F=7.38, df=1/111, p=0.008). These three scores were higher in subjects using haloperidol than in those without. In the same analysis we could not find any effects of age, TUD, education and familial history of psychosis. There were no significant effects of risperidone, clozapine, perphenazine, antidepressants, mood stabilizers and anxiolytics on the rating scores.

### Differences in TRYCAT ratios between the three WHO-QoL groups

Table 2, multivariate regression #3 shows that there was a significant association between the three ratios and WHO-QoL groups. **ESF Figure 3** shows the z-transformed values of the three ratios in the WHO-QoL groups, while table 3 shows the model-generated means. Table 3 shows that there were also significant differences in the TRYCAT LV scores extracted from the three ratios (univariate GLM analysis: F=9.29, df=2/114, p<0.001, R^2^=0.144). Post-hoc analyses show that the TRYCATs are significantly higher in the lower WHO-QoL groups as compared with the normal WHO-QoL group.

**Figure 3.**
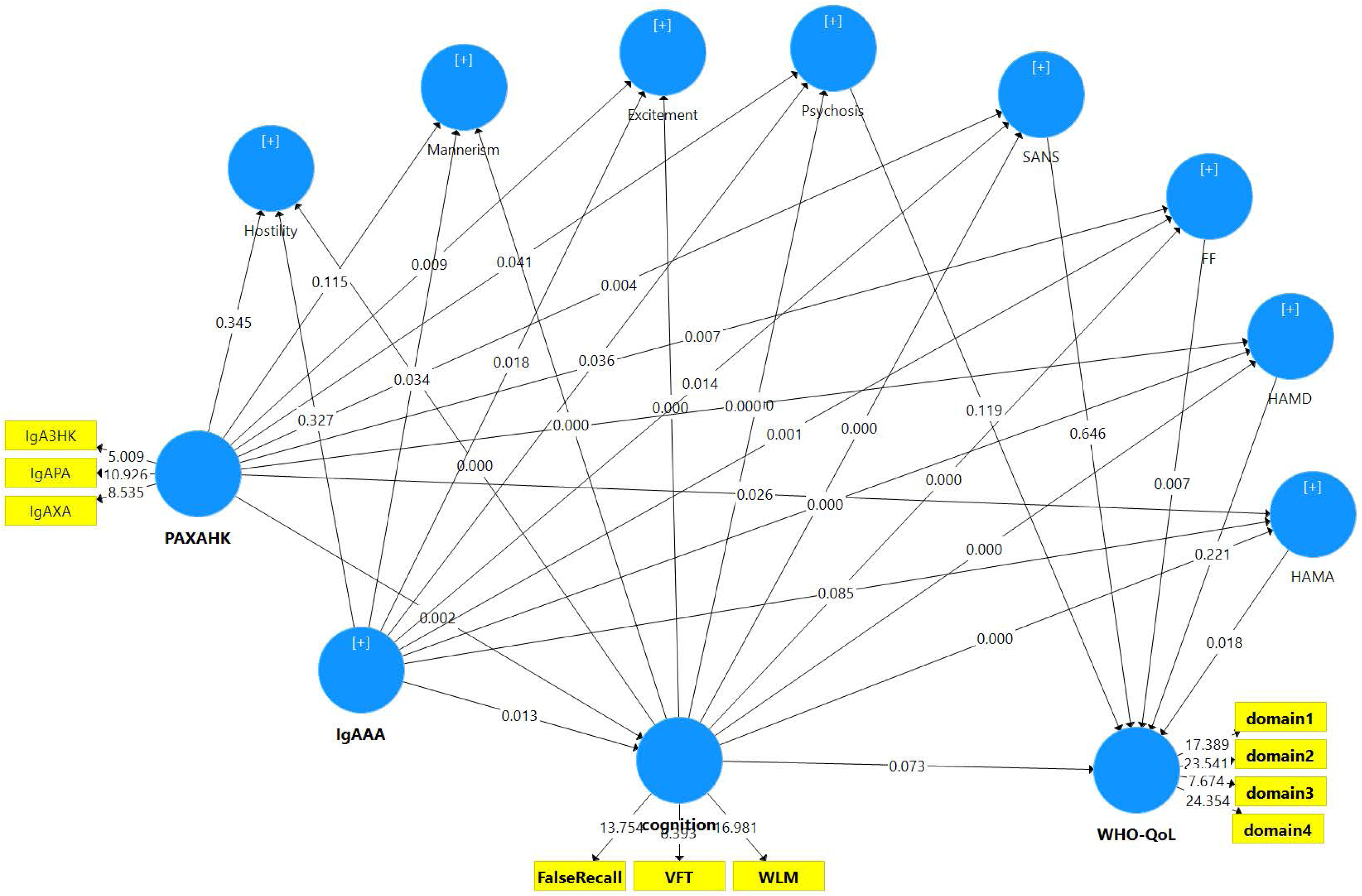
Results of Partial Least Squares path modeling analysis (with exact p-values) with the 4 domains of the WHO-QoL as indicator variables of the latent construct “WHO-QoL” and all symptom dimensions, cognitive tests and tryptophan catabolites, namely picolinic acid (PA), xanthurenic acd (XA), 3-OH-kynurenine (3HK) and anthranilic acid (AA) as input variables. See legends to ESF Figure 2, 3 and 4 for explained abbreviations.

### Differences in the 5 CERAD tests between the three WHO-QoL groups

Table 2, multivariate GLM regression #4 shows that there was a significant association between WHO-QoL groups and CERAD tests after adjusting for effects of education and sex. Table 3 also shows the CERAD LV scores obtained by univariate GLM analysis (F=27.73, df=4/113, p<0.001, R^2^=0.495) which yielded effects of the WHO-QoL groups (F=21.19, df=2/113, p<0.001) and education (F=30.55, df=1/113, p<0.001, R^2^=0.213), but not sex (F=3.34, df=1/113, p=0.070). **ESF Figure 4** displays the z-transformed values of the 5 CERAD tests, while Table 3 displays the model-generated estimated marginal mean values. WLM, False Recall and CERAD LV scores were significantly different between the three groups and VFT and MMSE scores were different between the normal WHO-QoL groups and lower WHO-QoL groups. Table 2 shows that there was also a significant effect of education and sex on the CERAD tests. Tests for between-subject effects showed significant effects of education on WLM (F=23.35, df=1/110, p<0.001), VFT (F=16.00, df=1/110, p<0.001), MMSE (F=64.02, df=1/110, p<0.001) and False Recall (F=10.85, df=1/110, p=0.002), but not WLRecog (F=3.78, df=1/110, p=0.055). After p-correction, there were also significant effects of sex on WLM (p=0.027), MMSE (p=0.035) (both lower in men) and False Recall (p=0.027, higher in men). There were no significant effects of the drug state of the patients on the CERAD tests, namely risperidone (p=0.943), clozapine (p=0.735), haloperidol (p=0.305), perphenazine (p=0.888), antidepressants (p=0.888), mood stabilizers (p=0.888) and anxiolytics (p=0.147).

### Results of multiple linear regression analyses (MLRA)

**Table 4** shows the results of 5 different MLRA with the WHO-QoL scores, either total or the 4 domain scores, as dependent variables and TRYCAT ratios, CERAD tests, PHEM and DAPS symptoms (and age, sex, education, diagnosis schizophrenia and deficit schizophrenia entered as dummy variables) as explaining variables. We found that 54.0% of the variance in the total WHO-QoL score was explained by HAMA, SANS and FF scores and VFT. Univariate analysis showed that 48.5% of the variance in the LV extracted from the 4 domains was explained by the general psychopathology LV extracted from the 8 PHEMN/DAPS symptoms (F=109.17, df=1/116, p<0.001) and that DAPS (F=106.58, df=1/116, p<0.001, R^2^=47.9%) had more impact than PHEMN (F=67.12, df=1/116, p<0.001, R^2^=36.7%) symptoms. 46.7% of the variance in Domain1 was explained by psychosis, HAMA, age and IgM KA_3HK ratio. Domain2 (48.6%) was best explained by FF, VFT and false recall. The best predictors of Domain3 (28.1%) were HAMA, hostility and VFT, whereas Domain4 was best predicted by FF, deficit schizophrenia, HAMA, VFT and the ΔNOX_PRO explaining 41.3% of the variance. The diagnosis of schizophrenia (versus controls) was not significant in these regressions.

**Table 4.** Results of stepwise multiple regression analyses with the total and domain scores of the World Health Organization Quality of Life instrument-Abbreviated version (WHO-QoL-BREF) as dependent variables and tryptophan catabolite (TRYCATs) ratios, Consortium to Establish a Registry for Alzheimer’s Disease (CERAD) tests, and symptom dimensions as primary explanatory variables.

| Dependent Variables | Significant explanatory variables | t | p | R<sup>2</sup> (%) | F | df | P |
| --- | --- | --- | --- | --- | --- | --- | --- |
| **WHO-QoL total** | FF<br>SANS<br>HAMA<br>VFT | -2.32<br>-2.38<br>-2.82<br>+2.41 | 0.022<br>0.019<br>0.006<br>0.017 | 52.6 | 31.03 | 4/112 | <0.001 |
| **Domain 1** | Psychosis<br>HAMA<br>IgM KA_3HK<br>Age | -4.37<br>-3.79<br>-2.89<br>-2.60 | <0.001<br><0.001<br>0.005<br>0.011 | 46.7 | 24.30 | 4/111 | <0.001 |
| **Domain 2** | FF<br>VFT<br>FalseRecall | -6.17<br>+2.72<br>-2.25 | <0.001<br>0.008<br>0.026 | 48.6 | 35.23 | 3/112 | <0.001 |
| **Domain 3** | HAMA<br>Hostility<br>VFT | -3.89<br>-2.43<br>+2.03 | <0.001<br>0.017<br>0.045 | 28.1 | 14.59 | 3/112 | <0.001 |
| **Domain 4** | FF<br>Deficit schizophrenia (Yes/No)<br>VFT<br>HAMA<br>ΔNOX_PRO | -2.84<br>-3.24<br>+2.35<br>-2.47<br>+2.05 | 0.005<br>0.002<br>0.021<br>0.015<br>0.043 | 51.3 | 23.20 | 5/110 | <0.001 |

HAMA: Hamilton Anxiety or Depression Rating Scale.
FF: the Fibromyalgia and Chronic Fatigue Syndrome Rating scale score
IgM KA\_3HK: computed as $z\text{IgM KA} - z3\text{HK}$ (index of lowered regulation of kynurenic acid versus 3-OH-kynurenine). $\Delta\text{NOX\_PRO}$ : computed as $z\text{IgA} (z\text{QA} + z\text{PA} + z\text{XA} + z3\text{HK} - z\text{AA} - z\text{KA}) - z\text{IgM} (z\text{QA} + z\text{PA} + z\text{XA} + 3\text{HK} - z\text{AA} - z\text{KA})$ (a more comprehensive index of increased noxious potential); QA: quinolinic acid; PA: picolinic acid; XA: xanthurenic acid; 3HK: 3-OH-kynurenine; AA: anthranilic acid; KA: kynurenic acid
VFT: verbal fluency test
Psychosis and Hostility: computed as z unit weighted composite scores using symptoms of the Brief Psychiatric Rating Scale and the Positive and Negative Syndrome Scale
SANS: total score on the Scale for the Assessment of Negative Symptoms

### Results of neural network analysis

Employing MLP NN we predicted the WHO-QoL score (output variable) using TRYCAT ratios, CERAD tests, PHEMN and DAPS symptoms (and age, sex, education) as input variables. Automatic architecture training of the network delineated the best model employing 2 hidden layers. Each layer contained 2 units, the first layer with hyperbolic tangent as activation function and the output layer with identity as activation factor, while using the sum of squares error function. **Table 5** shows the model summary of the trained Neural Network. The sum of squares error term is minimized during training, showing that the neural model has learnt to generalize from the trend. The training, testing and holdout samples showed an error term that was fairly constant with a lower error term in the testing and holdout samples than in the training sample, indicating that the model is not overtrained. The Spearman correlation coefficient between the WHO-QoL score and the model-predicted value is 0.744 (p<0.001, n=113). **ESF Figure 5** shows the relevance chart displaying relative and normalized importances of all input variables. The FF score was the most important determinant of the predictive power of the model, followed at a distance by HAMA, SANS and MMSE and again at a distance by hostility, the IgA NOX_PRO ratio and HAMD.

**Table 5.** Results on Neural Network analysis with WHO-QoL total score as dependent variable

|  |  |  |
| --- | --- | --- |
| Training | Sum of squares error | 13.878 |
|  | Relative error | 0.544 |
| Testing | Sum of squares error | 4.860 |
|  | Relative error | 0.368 |
| Hold-out | Relative error | 0.431 |
| Correlation | WHO-QoL total score<br>and predicted score | r=0.744<br>p<0.001 |

### Results of PLS-SEM path modeling

In model 1 (**Figure 1**) we examine the associations between the 4 WHO-QoL domains entered as indicator variables (in a reflective model) as final LV response variable. Symptoms (all 8 symptoms as indicators of the latent construct “general psychopathology”), cognition (5 CERAD tests as indicators of “cognitive function”) and all 3 TRYCAT ratios (as indicators of “TRYCAT patterning”) were used as direct input variables (all reflective), whereas TRYCATs and cognition also served as predictors of symptoms, and TRYCAT ratios as predictors of cognitive functions. Mannerism, MMSE and WLRecog were removed because they showed loadings <0.500 on their factors. The overall model fit was good (SRMR=0.073), and the construct reliability and discriminant validity of the 4 latent constructs were good to excellent (all Cronbach’s alpha > 0.8, composite reliability > 0.8 and average variance extracted > 0.6). The construct crossvalidated redundancies and communalities were adequate. Finally, all factor loadings of the 4 constructs showed loadings > 0.55 (all p<0.001), indicating that the convergent validity is adequate. Figure 6 shows the path coefficients with exact p value for all causal links. 64.4% of the variance in the 4 QoL domains was explained by general psychopathology, whereas cognitive impairments and TRYCATs patterning were not significant. 64.4% of the variance in general psychopathology was explained by cognitive impairments and TRYCAT ratios, while 12.2% of the variance in cognitive impairments was explained by TRYCAT ratios. Moreover, there were specific indirect effects of TRYCAT ratios on general psychopathology mediated by cognition (t=+3.84, p<0.001), and on WHO-QoL domains mediated by symptoms (t=-2.00, p=0.046) and the path from cognitive disorders to symptoms (t=-3.00, p=0.003). Total effects showed significant effects of TRYCATs (t=-2.79, p=0.005) and cognitive functions (t=+8.47, p<0.001) on the 4 QoL domains.

In a second model (**Figure 2**) we aimed to decipher the associations between the 4 domains (each entered as a separate outcome variable) and PHEMN and DAPS (each represented by LVs extracted from the 5 PHEMN and 3 DAPS scores, respectively), while TRYCATs and CERAD were entered as input variables. In order to examine the indirect effects of sex and education on WHO-QoL domains we entered sex and education as direct predictors of the CERAD LV, while also TRYCATs predicted cognitive tests. Again, all model quality data were good, including model fit SRMR=0.054. Figure 7 shows the path coefficients with exact p values for all causal links. We found that 39.9% of the variance in Domain1 was explained by DAPS and PHEMN, and 46.8% of the variance in Domain2, 20.9% in Domain3 and 46.0% in Domain4 by DAPS only. The TRYCAT LV, sex and education explained 50.9% of the variance in the CERAD LV. There were total direct effects of CERAD LV, education and TRYCATs (all at p<0.001) and sex (all <0.01) on Domain 1, 2, 3 and 4 scores and on DAPS and PHEMN symptoms (all p<0.001). There were specific indirect effects of TRYCATs on Domain1 mediated by DAPS (t=-2.33, p=0.020) and the path from CERAD to DAPS (t=-3.13, p=0.002) and from CERAD to PHEMN (t=-2.28, p=0.023). There were also specific indirect effects of TRYCATs on Domain2 mediated by DAPS (t=-2.24, p=0.025) and the path from CERAD to DAPS (t=-3.76, p<0.001); on Domain3 mediated by the path from CERAD to DAPS (t=-2.08, p=0.038); and on Domain4 mediated by DAPS (t=-2.35, p=0.019) and the path from CERAD to DAPS (t=-3.65, p<0.001). There were significant specific indirect effects of male sex on DAPS (t=+2.54, p=0.011) and PHEMN (t=+2.50, p=0.013) mediated by the CERAD LV. There were significant specific indirect effects of sex on Domain1 (t=-2.04, p=0.042), Domain2 (t=-2.34, p=0.019) and Domain4 (t=-2.21, p=0.027) mediated via effects of CERAD LV on DAPS symptoms. There were significant specific indirect effects of education on DAPS (t=-5.79, p<0.001) and PHEMN (t=-6.67, p<0.001) mediated by the CERAD LV. There were significant specific indirect effects of education on Domain1 (t=+3.04, p=0.002), Domain2 (t=+3.76, p<0.001), Domain3 (t=+1.96, p=0.048) and Domain4 (t=++3.78, p<0.001) mediated by effects of CERAD LV on DAPS symptoms, and on Domain1 (t=+2.51, p=0.012) mediated by effects of CERAD LV on PHEMN symptoms.

In the third model (**Figure 3**) we examine the associations between the latent construct WHO-QoL with the 4 domain scores entered as indicator variables, and all symptoms entered as separate variables, while CERAD tests were entered as a latent construct. In addition, the noxious (XA, QA, PA and 3HK) and generally more protective (AA and KA) TRYCATs were entered as indicators of two latent constructs, namely noxious versus protective TRYCATs. Two CERAD tests, QA and KA were removed from the latent constructs because the factor loadings did not reach the required criteria. After this, the model quality data complied with the quality criteria with a general model fit SRMR=0.035. Path analysis showed that 58.2% of the variance in the WHO-QoL LV was explained by FF and HAMA (the other symptoms were non-significant). We found that 26.5% of the variance in HAMA, 61.6% in HAMD, 43.9% in FF, 67.2% in SANS, 52.0% in excitement and 43.2% in psychosis was explained the regression on CERAD LV (inversely), the noxious TRYCAT LV (positively) and AA (inversely). 26.5% of the variance in hostility was explained by CERAD LV, whilst 36.6% of the variance in mannerism was explained by CERAD LV and AA (both inversely). 15.8% in the LV CERAD score was explained by noxious TRYCAT LV (inversely) and AA (positively). There were significant total effects of noxious TRYCAT LV (t=-3.72, p=0.001), AA (t=+3.24, p=0.001), CERAD LV (t=+6.10, p<0.001), FF (t=-2.58, p=0.010) and HAMA (t=-2.21, p=0.027) on the WHO-QoL LV.

## Discussion

The first major finding of this study is that lowered HR-QoL in schizophrenia is strongly predicted by clinical symptoms, whereby general psychopathology explains around 48.5% of the variance. These findings extend previous knowledge that psychiatric symptoms in general, and positive, negative and mood symptoms may impact HR-QoL in schizophrenia (Galuppi et al., 2010; Sim et al., 2004; Ritsner et al., 2005; Becker et al., 2005; Fitzgerald et al., 2001; Gorna et al., 2014; Savill et al., 2016). In accordance with Sum et al. (2018) we found that deficit schizophrenia was accompanied by significantly lowered HR-QoL as compared with nondeficit schizophrenia. This association may be explained since the deficit phenotype is characterized by greater severity of illness as compared with nondeficit schizophrenia (Kanchanatawan et al., 2018d). Nevertheless, the current study found that severity of overall symptomatology had a stronger impact on HR-QoL than the deficit phenotype per se.

Previously we showed that schizophrenia symptomatology comprises two inter-related dimensions, namely DAPS and PHEMN (Kanchanatawan et al., 2018e). Importantly, the current study found that DAPS symptoms have a much stronger impact on HR-QoL than PHEMN symptoms. As reviewed in the introduction, previous research often yielded contradictory results with some studies reporting that depressive, but not negative or positive, symptoms predict HR-QoL, while other studies showed that negative as well as positive symptoms predict HR-QoL (Becker et al., 2005; Fitzgerald et al., 2001; Norman et al., 2000; Savill et al., 2016). Some of the discrepancies in previous studies may be explained by our findings that mood, negative and positive symptoms are strongly interrelated and that positive symptoms should be further “dissected” into the more relevant PHEM symptoms (Kanchanatawan et al., 2018e). Moreover when we examined all 8 PHEMN and DAPS symptoms separately, we detected that the lowered HR-QoL in schizophrenia is best predicted by severity of physio-somatic and anxiety symptoms, although also psychotic and hostility symptoms impacted domains 1 and 3 of the HR-QoL, respectively. To the best of our knowledge this is a first study showing that physio-somatic symptoms have a stronger impact on HR-QoL than psychotic, negative and depressive symptoms. Nevertheless, inspecting the different questions in the HR-QoL reveals that many items reflect physio-somatic symptoms. For example, Domain1 (physical health) comprises items such as energy and fatigue, pain, sleep and rest and work capacity, all items that are affected or reflect physio-somatic symptoms. Domain2 (psychological) comprises items such as memory and concentration, negative feelings and self-esteem and Domain3 (social relationships) comprises sexual activity, which all belong to the physio-somatic dimension. Interestingly, a number of studies have shown that patients with chronic fatigue syndrome, a medical illness characterized by physio-somatic symptoms, show a lowered HR-QoL (Schweitzer et al., 1995; Hvidberg et al., 2015; Van Heck and de Vries, 2002). These studies indicate that chronic fatigue syndrome has a HR-QoL burden affecting a wide range of factors and especially social functions, findings which could explain why in schizophrenia the FF score strongly impacts Domain4.

The second major finding of this study is that memory impairments strongly impact HR-QoL and that these effects are mediated via DAPS/PHEMN symptoms. Also, previous studies showed that cognitive deficits in schizophrenia may impact HR-QoL. For example, Ueoka et al. (2010) reported that not only depressive and negative symptoms impact HR-QoL, but also cognitive tests including verbal memory, attention and speed of information processing. Alptekin et al. (2005) found that executive dysfunctions and impairments in working memory have direct effects on HR-QoL, especially in the social domains. A meta-analysis (Tolman and Kurtz, 2012) examined 20 HR-QoL studies with 1615 patients, 10 using objective (clinician rated) and 10 subjective (patient satisfaction) QoL measurements. The objective analyses showed small–moderate associations between executive functions, working memory, verbal ability, verbal list learning, and processing speed and HR-QOL, whereas the subjective analyses showed nonsignificant or even inverse relationships (Tolman and Kurtz, 2012). Mohamed et al. (2008) reported that not only positive and negative symptoms but also cognitive deficits are associated with HR-QoL in schizophrenia, although symptoms contributed more to HR-QoL than cognitive impairments and that the impact of positive symptoms was about equal to the effect of cognition. Keefe and Harvey (2012) reviewed that cognitive impairments in schizophrenia are associated with measures of community functioning, including independent living, functional capacity, employment, work behavior and performance. For example, cognitive performance may have a stronger impact on the ability to work than clinical symptoms (McGurk and Mueser, 2003).

Our results support previous theories that cognitive impairments, including learning, attention, memory and executive functions, may generate false memories and psychosis and consequently underpin schizophrenia symptomatology (Orellana and Slachevsky, 2013; Keefe and Harvey, 2012; Corlett et al., 2007). For example, verbal memory deficits and attentional impairments in ultra-high risk individuals predict psychotic symptoms (Brewer et al., 2005; Hawkins et al., 2004). Given that cognitive deficits precede the onset of psychosis and that cognitive deficits are a result of central neuro-circuitry dysfunctions, which precede psychotic symptoms (Harvey et al., 2006; Tamminga et al., 2006), it is safe to conclude that cognitive deficits contribute to the clinical symptoms of schizophrenia and subsequently to a lowered HR-QoL. Likewise, it is probable that DAPS symptoms, which are strongly related to PHEMN symptoms, may be a consequence of cognitive deficits. For example, false memory creation may be present prior to subsequent depression and therefore may be a risk factor for depressive symptoms (Myhre, 2015).

The third major finding of our study is that changes in TRYCAT patterning, including increased levels of noxious TRYCATS (PA, XA and 3HK) and lowered levels of protective (AA) TRYCATs, significantly impact HR-QoL and that these effects are mediated via cognitive impairments and DAPS/PHEMN symptoms. As such a large part of the variance (> 50%) in HR-QoL measurements can be explained by direct and indirect effects of TRYCAT patterning, memory deficits and the DAPS/PHEMN symptoms combined. Previously, we have reviewed the many pathways by which noxious TRYCATs may cause neurocognitive impairments and clinical symptoms. Briefly: a) the TRYCAT pathway is induced by M1 macrophage cytokines (increased interleukin-1, interleukin-6 and tumor necrosis factor-α production), T helper-1 cytokines (increased interferon-γ production), and reactive oxygen species (ROS), which have neurotoxic and cytotoxic effects thereby causing neuroprogressive processes; and b) noxious TRYCATS may exert excitotoxic, neurotoxic and cytotoxic effects and additionally cause oxidative, nitrosative and inflammatory responses which all may cause neuroprogression thereby contributing to cognitive impairments and clinical symptoms (Kanchanatawan et al., 2018a; 2018c).

Accordingly, we propose that previous theories concerning the role of cognitive impairments (Orellana and Slachevsky, 2013; Keefe and Harvey, 2012) and other psychological factors including awareness (Frith, 2000) should be reformulated. Thus, the more distal outcome of schizophrenia (lowered HR-QoL) and general and PHEMN / DAPS symptom dimensions are affected by a broad range of cognitive impairments (including episodic and semantic memory, executive functions and sustained attention and emotion recognition), which in turn are associated with activated neuro-immune pathways, while the effects of the latter on the outcome data are mediated by cognitive deficits and their effects on DAPS, physio-somatic and anxiety symptoms.

Finally, the current study shows that the reduction in CERAD performance may be explained not only by changes in TRYCAT patterning, but also by male sex and lower education. Similar effects were reported by Carduso et al. (2005) who found that male gender is accompanied by lowered HR-QoL. Nevertheless, here we report that the effects of male sex on HR-QoL are mediated by on memory and their effects on clinical symptoms. In this respect, it is interesting to note that schizophrenia has a higher prevalence and worse outcome in males (Leung and Chue, 2000; Häfner, 2003). It is possible that these sex effects are related to effects of oestrogens, which may improve cognitive functions by enhancing spinogenesis and synaptogenesis in the hippocampus (Häfner, 2003). The results of the current study also show that higher education protects against a worse intermediate and distal outcome (lowered HR-QoL) of schizophrenia. Previous studies also showed that schizophrenia patients with higher education have higher HR-QoL as compared with those with lower education (Carduso et al., 2005; Choo et al., 2017). Nevertheless, here we found that the effects of education on HR-QoL are mediated by memory deficits and their effects on DAPS (Domain 1, 2, 3 and 4) and PHEMN (Domain 1 only) symptoms. These protective effects of higher education may be explained by a) better cognitive reserve and thus increased resilience to the detrimental effects of brain damage (Richards and Sacker, 2003), and b) increased protection against ROS and lipid peroxidation (Maes et al., 2018), which may be explained by a healthier lifestyle, better nutrition and more exercise in more educated subjects (Maes et al., 2018; Lobo et al., 2010; Moylan et al., 2013).

In conclusion, lowered HR-Qol in schizophrenia is to a great extent predicted by changes in TRYCAT patterning, impairments in memory (episodic and semantic), clinical symptoms especially DAPS and physio-somatic symptoms. Noxious (PA, XA, 3HK) and protective (AA) TRYCATs strongly impact HR-QoL and the effects of noxious/protective TRYCATs are mediated by memory impairments and the effects of the latter on clinical symptoms.

## Acknowledgement

None

## Contributors

MM and BK designed the study. BK recruited patients and completed diagnostic interviews and rating scales measurements. MM carried out statistical analyses Both BK and MM contributed to interpretation of the data and writing of the manuscript.

